# Atypical Chemokine Receptor 3 ‘Senses’ CXC Chemokine Receptor 4 Activation Through GPCR Kinase Phosphorylation

**DOI:** 10.1101/2023.02.25.530029

**Authors:** Christopher T. Schafer, Qiuyan Chen, John J. G. Tesmer, Tracy M. Handel

**Affiliations:** Skaggs School of Pharmacy and Pharmaceutical Sciences, University of California, San Diego, La Jolla, USA; Departments of Biological Sciences and of Medicinal Chemistry and Molecular Pharmacology, Purdue University, West Lafayette, IN, USA; Dept. of Biochemistry and Molecular Biology, Indiana University School of Medicine, Indianapolis, IN, USA

## Abstract

Atypical chemokine receptor 3 (ACKR3) is an arrestin-biased receptor that regulates extracellular chemokine levels through scavenging. The scavenging action mediates the availability of the chemokine CXCL12 for the G protein-coupled receptor (GPCR) CXCR4 and requires phosphorylation of the ACKR3 C-terminus by GPCR kinases (GRKs). ACKR3 is phosphorylated by GRK2 and GRK5, but the mechanisms by which these kinases regulate the receptor are unresolved. Here we mapped the phosphorylation patterns and determined that GRK5 phosphorylation of ACKR3 dominates β-arrestin recruitment and chemokine scavenging over GRK2. Co-activation of CXCR4 significantly enhanced phosphorylation by GRK2 through the liberation of Gβγ. These results suggest that ACKR3 ‘senses’ CXCR4 activation through a GRK2-dependent crosstalk mechanism. Surprisingly, we also found that despite the requirement for phosphorylation, and the fact that most ligands promote β-arrestin recruitment, β-arrestins are dispensable for ACKR3 internalization and scavenging, suggesting a yet to be determined function for these adapter proteins.

## Introduction

Chemokines receptors mediate cell migration in the context of immune system function, development, and disease by responding to small (8-14 kDa) protein agonists (chemokines) and activating G protein signaling cascades that lead to cell motility^1^. Cell positioning also depends on the establishment of localized chemokine gradients, which provide directional cues for migrating cells^2, 3^. Thus, in addition to G protein-coupled, canonical chemokine receptors (CCKRs) that directly mediate cell movement, a subclass of G protein-independent, “atypical” chemokine receptors (ACKRs) indirectly contribute to migration by controlling extracellular chemokine concentrations and shaping chemokine gradients through ’scavenging.’ Scavenging by ACKRs restricts CCKR access to chemokines and regulates canonical receptor down regulation, thereby maintaining chemokine receptor responsiveness and ability to promote cell migration^4, 5^.

ACKR3 (also known as CXCR7) is an atypical receptor that shares the endogenous ligands of the CCKRs: CXCL11 (an agonist of CXCR3) and CXCL12 (the sole agonist of CXCR4)^6–8^. Although the receptor has been reported to function on its own^9, 10^, ACKR3 also cooperates with CXCR3 and CXCR4, in part by scavenging CXCL11 and CXCL12^7^. Scavenging involves internalization of receptor-bound chemokines into the cell for degradation, followed by recycling of the receptor back to the cell surface for further rounds of chemokine consumption^7, 8^. The importance of ACKR3 is underscored by the fact that its absence leads to severe defects in CXCR4-mediated interneuron migration^11^ and development of the zebrafish lateral line primordium^12^.

Like other ACKRs, ACKR3 does not activate heterotrimeric G proteins^11, 13^, with the exception of cell-specific coupling in astrocytes^14, 15^. However, it is phosphorylated by GPCR kinases (GRKs), which results in the recruitment of β-arrestins^9, 11, 13, 16^. In some studies, β-arrestin2 has been reported to be required for scavenging^4, 17^, yet others show efficient chemokine uptake in cells lacking β-arrestins^11, 18^. Nevertheless, phosphorylation by GRKs seems critical for ACKR3 scavenging^11, 16^. C-terminal phosphorylation is increased upon CXCL12 stimulation of ACRK3 and switches the receptor from ubiquitin-mediated lysosomal degradation towards plasma membrane recycling^19, 20^.

Most studies have implicated GRK2 in the internalization of ACKR3 as well as ACKR3-mediated arrestin recruitment and chemokine scavenging^11, 16^. Although GRK5 also phosphorylates ACKR3^11^, the importance of GRK5 and the relative contribution and precise roles of GRK2 vs. GRK5 have not been elucidated. Here we compare the impact of GRK2 and GRK5 on ACKR3 internalization, β arrestin recruitment, and scavenging. Although published studies have focused on GRK2 as the key ACKR3 kinase in HEK293 cells and primary neurons^11, 16^, our results suggest that GRK5 dominates ACKR3 phosphorylation in HEK293A cells. However, we also find that phosphorylation of ACKR3 by GRK2 is increased by co-activation of CXCR4. The data are consistent with the dependence of GRK2 on Gβγ^21^, which is liberated by CXCL12 stimulation of CXCR4 but not by ACKR3, and with the Gβγ independence of GRK5^22^. These results suggest a kinase-mediated mechanism that enables ACKR3 to sense the activation of CXCR4 and respond in a cell-specific manner depending on the co-expression and activation of the canonical GPCR.

## Results

### GRKs are necessary for efficient scavenging of CXCL12 by ACKR3

Although GRK2 has been shown to contribute to ACKR3 internalization, β-arrestin recruitment, and chemokine scavenging, the relative importance of GRK2 versus other GRKs in these processes has not been determined^11, 16^. To assess the impact of GRKs on these ACKR3 functions, we first tested ACKR3 internalization and chemokine scavenging in HEK293A cells in which GRKs 2, 3, 5, and 6 were knocked out by CRISPR (ΔGRK2/3/5/6)^23^. ACKR3 internalization was monitored by bystander bioluminescence resonance energy transfer (BRET) between ACKR3 C-terminally tagged with luciferase (ACKR3_RlucII) and GFP fused to a CAAX domain (rGFP_CAAX) that anchors it to the plasma membrane^24^. In wild-type (WT) cells, CXCL12 induced a rapid decrease (>35%) in BRET signal corresponding to ACKR3 internalization (Fig. 1A). The internalization was almost eliminated in ΔGRK2/3/5/6 cells, with only a 2% decrease in the BRET ratio, indicating a critical role for GRKs.

**Figure 1:**
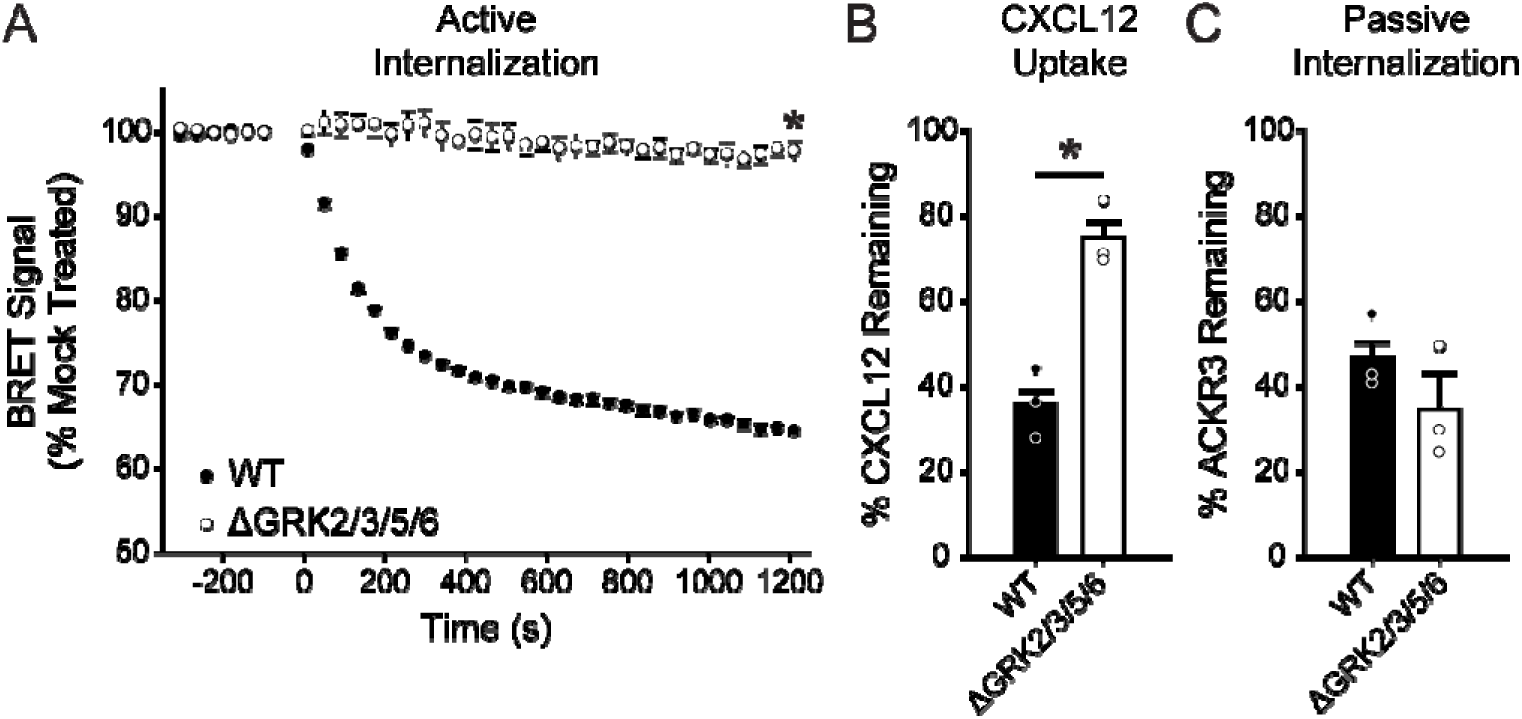
GRKs mediate efficient CXCL12-induced internalization and chemokine scavenging by ACKR3. A) CXCL12-promoted, active internalization following stimulation with 100 nM chemokine at 37 °C was monitored by BRET between ACKR3_RlucII and rGFP_CAAX in WT and ΔGRK2/3/5/6 HEK293A cells. B) Chemokine uptake by WT and ΔGRK2/3/5/6 cells expressing ACKR3 was detected by quantification of the remaining CXCL12 in the media by ELISA. The extent of scavenging was determined by comparison to cells transfected with empty vector. C) Constitutive cycling, or passive internalization, of ACKR3 in WT and ΔGRK2/3/5/6 cells was quantified by tracking the loss of ‘pre-labeled’ receptor by flow cytometry. Errors are reported as standard deviations and values are the average of three independent experiments performed in triplicate. Individual experiments are presented as points in B and C. Statistical significance was determined by one-way ANOVA followed by Bonferroni test. *P<0.001.

We next determined the effect of the ΔGRK2/3/5/6 knockout on ACKR3 scavenging by ELISA quantification of CXCL12 remaining in the bulk media following incubation with cells transfected with receptor or empty vector. Comparable receptor expression in the various cell lines was confirmed by luminescence from the C-terminal luciferase tag (Supplementary Fig. 1). In WT cells, ACKR3 efficiently removed ∼65% of CXCL12 in comparison to non-ACKR3 expressing cells after overnight incubation (Fig. 1B). In the absence of GRKs, the chemokine uptake was severely impaired, with about 80% remaining in the bulk media. Interestingly, it appears that ACRK3 retains the capacity to clear approximately 20% of the added CXCL12 (Fig 1B) even when internalization, as detected by BRET, is nearly eliminated (Fig. 1A). One possible explanation is that ACKR3 constitutively internalizes and recycles back to the plasma membrane, which has been previously shown to contribute to CXCL12 scavenging^13, 17^. Constitutive internalization is not detected by BRET, which is only sensitive to changes following a perturbation of the equilibrium state (such as ligand-induced translocation of the receptor from the plasma membrane to the inside of the cell). Thus, to ascertain the impact of GRKs on ACKR3 constitutive internalization and recycling, we employed a “pre-label” flow cytometry experiment^17^ instead of the BRET assay. In this experiment, surface receptors are first labeled with a non-conjugated anti-ACKR3 antibody at 4 °C (a temperature that halts internalization), and then warmed to 37 °C to allow for constitutive internalization and endocytic trafficking. The remaining original (pre-labeled) receptor is detected by fluorescent secondary antibody staining, and the level of constitutive internalization quantified by comparing the cells incubated at 37 °C with control samples held at 4 °C. In contrast to the ligand-stimulated internalization of the receptor detected by BRET, no significant difference between WT and ΔGRK2/3/5/6 cells was observed (Fig. 1C) consistent with constitutive trafficking occurring in the absence of GRKs. Note that in the remainder of the text, we interchangeably refer to BRET-detected internalization as “active internalization”, which as demonstrated here requires GRK-mediated phosphorylation, and constitutive internalization as “passive internalization”, which does not.

### GRK2 and GRK5 differentially regulate CXCL12-mediated β-arrestin recruitment and internalization by ACKR3

Having established the importance of GRKs to ACKR3 function, we next evaluated the relative contributions of specific GRKs. Using GFP10-tagged β-arrestin2 (GFP_βarr2) and ACKR3_RlucII, β-arrestin2 recruitment to ACKR3 was monitored by BRET in GRK2/3 and GRK5/6 CRISPR-knockout cells (ΔGRK2/3 and ΔGRK5/6 cells, respectively)^23^. Recruitment in ΔGRK5/6 cells was reduced to approximately 40% of the WT maximum (Fig. 2A). In contrast, recruitment in ΔGRK2/3 cells had an insignificant effect on the recruitment E_max_. As expected, β-arrestin recruitment in ΔGRK2/3/5/6 cells was nearly abolished. Prior experiments individually over-expressing the four GRKs showed that only

**Figure 2:**
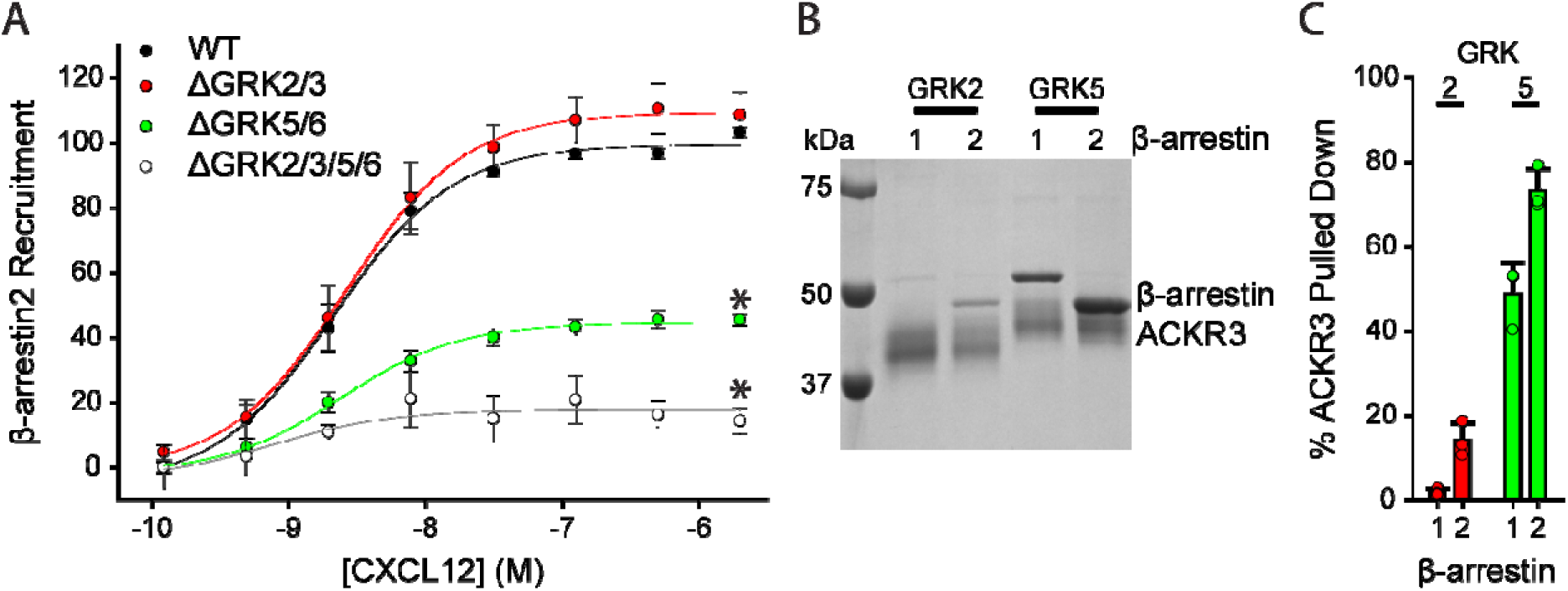
GRK5 plays a dominant role in ACKR3 phosphorylation and β-arrestin recruitment in HEK293A cells and with purified components. A) Recruitment of GFP_ βarr2 to ACKR3_RlucII observed by BRET in ΔGRK HEK293A cell lines across a titration of CXCL12 concentrations. Responses are normalized to WT ACKR3 and are a composite of three independent experiments performed in triplicate. B) Pulldowns of purified β-arrestin1 and 2 by purified ACKR3 phosphorylated in vitro by either GRK2 or GRK5. C) Quantification of the amount of β-arrestin pulled down from three independent experiments. The amount of β-arrestin pulled down in panel C is presented as a percentage of the ACKR3 band density from the same experiment. Errors are reported as standard deviation and significance in panel A is determined by extra sum of squares F-test, *P<0.001. EC_50_ values were not significantly different between the tested conditions.

GRK2 and GRK5 phosphorylate ACKR3^11^ and therefore we attribute the results using the ΔGRK2/3 and ΔGRK5/6 cells to the loss of GRK2 and GRK5, respectively. These results were corroborated by *in vitro* pulldowns of purified ACKR3-β-arrestin complexes phosphorylated with each kinase (Fig. 2B). The more extensive upward shift of the receptor band for GRK5-phosphorylated ACKR3 suggests greater phosphorylation compared to GRK2 under similar conditions. Additionally, the pulldowns of GRK5 phosphorylated ACKR3, revealed that approximately 50% and 70% of the receptor was complexed with β-arrestin1 and 2, respectively (Fig. 2C), whereas only 20% of the receptor was complexed with β-arrestin2 and almost no β-arrestin1 was detected in the pulldowns of GRK2 phosphorylated ACKR3. Together, the in cell and *in vitro* methods point to the same conclusion: that β-arrestin recruitment to ACKR3 is dominated by GRK5 phosphorylation under our conditions.

Because GRKs are necessary for CXCL12-mediated internalization of ACKR3, we also tested the impact of GRK2/3 or GRK5/6 phosphorylation on ligand-induced membrane trafficking. ACKR3 on the plasma membrane was once again monitored by BRET between ACKR3_RlucII and rGFP_CAAX. Similar to β-arrestin recruitment, active internalization of ACKR3 in ΔGRK5/6 cells was significantly reduced (∼25% BRET loss) compared to WT cells (∼35% BRET loss) (Fig. 3A). In contrast, the internalization was slightly enhanced (∼5%) in the ΔGRK2/3 cells. Together with the lack of effect on β-arrestin recruitment, this suggests little impact of GRK2 phosphorylation when GRK5 is present in these cells.

**Figure 3:**
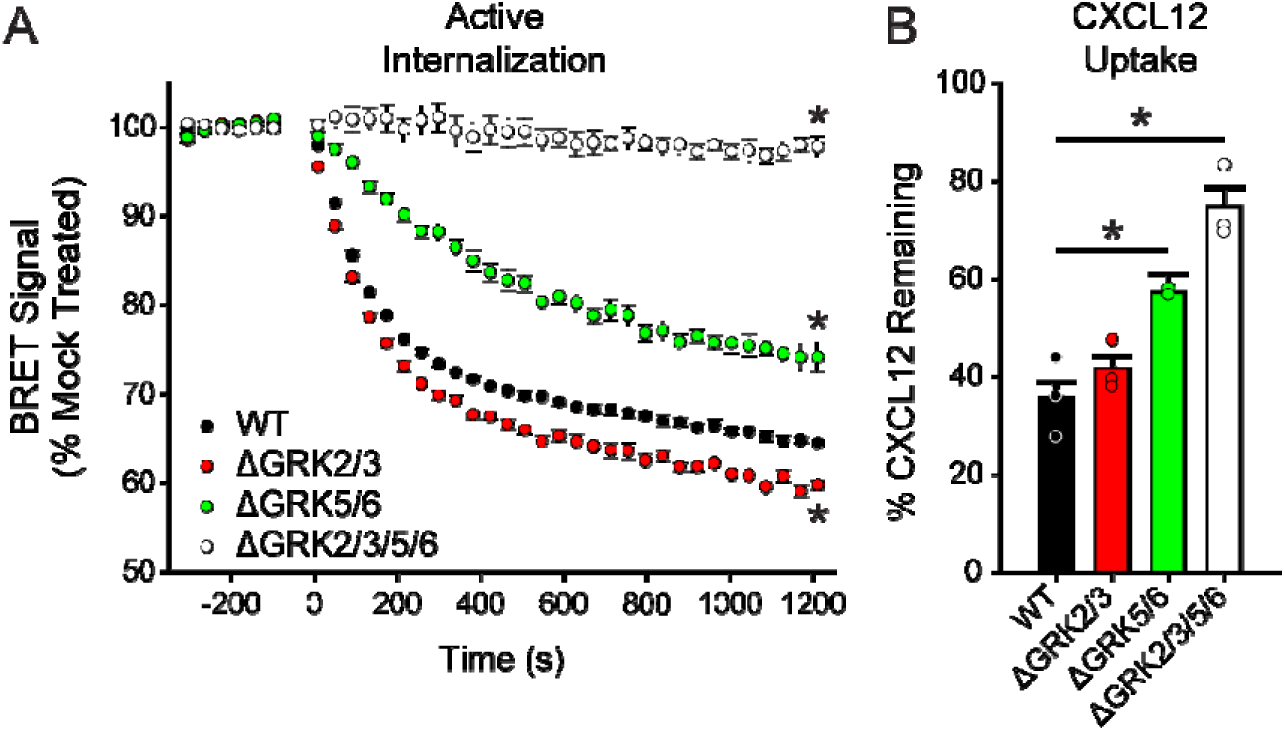
ACKR3 requires GRK5 for efficient active internalization and CXCL12 scavenging in HEK293A cells. A) Active, CXCL12-promoted internalization of ACKR3 monitored by BRET following stimulation with 100 nM CXCL12 in ΔGRK HEK293A cell lines at 37 °C. B) CXCL12 uptake by ACKR3 expressed in ΔGRK cell lines quantified by ELISA and presented as a percentage of identical cells transfected with empty vector. Data are composites of three independent experiments measured in triplicate and errors are reported as standard deviations and statistical significance was determined by one-way ANOVA followed by Bonferroni Test. *P<0.001. Data for WT and ΔGRK2/3/5/6 cells are repeated from Fig. 1 for comparison.

Finally, we tested how specific GRKs alter CXCL12 scavenging. ACKR3 only removed approximately 40% of CXCL12 when expressed in ΔGRK5/6 cells compared to more than 60% in WT cells. In contrast, CXCL12 scavenging in ΔGRK2/3 cells was indistinguishable from WT (Fig. 3B), again consistent with little contribution of GRK2 when GRK5 is present. However, the ΔGRK2/3/5/6 cells showed even less scavenging than the ΔGRK5/6 cells (∼20% scavenged). Moreover, all tested ACKR3 functions showed a greater decrease in the ΔGRK2/3/5/6 cells compared to the ΔGRK5/6 cells, suggesting that GRK2 and 5 may synergize.

### Mutation of ACKR3 phosphorylation site clusters correlate with effects of GRK knockout cells

The different functional responses of GRK2 and GRK5 phosphorylated ACKR3 suggest that the kinases introduce distinct phosphorylation barcodes and/or levels of phosphorylation. Accordingly, mass spectrometry of purified and *in vitro* phosphorylated ACKR3 was used to identify the positions modified by each kinase (Supplementary Fig 2). As shown in Fig. 4A, both GRK2 and GRK5 preferentially phosphorylate distal positions of the receptor C-terminus (S347/S350/T352/S355), that were previously shown to be important for β-arrestin binding to ACKR3^16^ and efficient chemokine scavenging^13^. GRK5 also preferentially phosphorylated proximal sites (S335/T338/T341), while GRK2 phosphorylated terminal positions (S360/T361). To further explore the role of these sites, we mutated to alanine all Ser/Thr residues in each of the three clusters as indicated in Fig. 4A.

**Figure 4:**
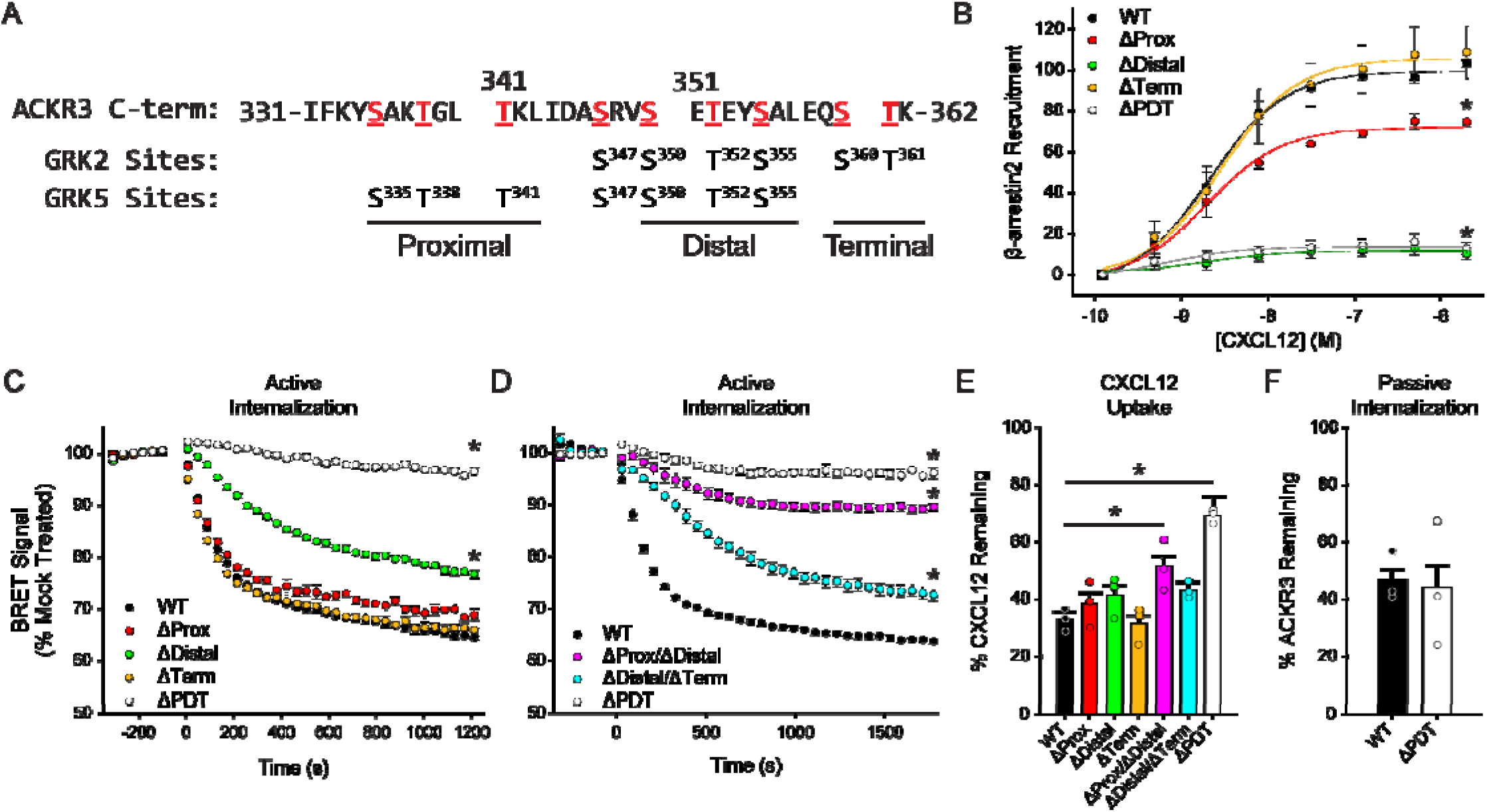
Specific phosphorylation motifs differently contribute to CXCL12 responses by ACKR3 in HEK293A cells. A) ACKR3 was phosphorylated by either GRK2 or GRK5 in vitro and the specific sites of modification were determined by mass spectrometry. The detected phosphorylation sites are highlighted in red. Phosphorylated positions in the ACKR3 C-terminus were divided into three clusters and replaced by alanine to produce ΔProx, ΔDistal, ΔTerm, and ΔPDT receptor constructs. B) Recruitment of GFP_βarr2 to phosphorylation deficient ACKR3 constructs expressed in WT HEK293A cells and tested across a titration of CXCL12 and detected by BRET. Values represent three independent experiments performed in triplicate and normalized to WT ACKR3 recruitment. C,D) Active internalization of individual (C) and multiple (D) phosphorylation cluster substitutions tracked by the loss of BRET between ACKR3_RlucII and rGFP_CAAX after CXCL12 addition. E) CXCL12 uptake by phosphorylation deficient ACKR3 constructs measured by remaining chemokine and compared to cells transfected with empty vector. F) Passive, agonist-independent internalization of the triple phosphorylation substitution observed by the pre-label flow cytometry experiment. WT ACKR3 data in B, C, E, and F was repeated from Fig. 1 and Fig. 2A for comparison. All errors bars represent standard deviations and statistical significance was determined by (B) extra sum of squares F-test and (C,D,E) one-way ANOVA followed by Bonferroni test. *P<0.001.

Mutation of the distal sites (ΔDistal) completely eliminated β-arrestin2 recruitment (Fig. 4B) that the distal mutation had a greater impact on recruitment than loss of GRK5/6 or GRK2/3, even though both GRKs phosphorylate the distal sites, is likely due to compensation by the remaining kinases in the knockout cells. Mutation of the terminal phosphate sites (ΔTerm), a motif specific for GRK2, displayed similar β-arrestin recruitment to WT ACKR3 and replicated the limited effect observed in ΔGRK2/3 cells. Partial impairment exhibited by the proximal site mutant (ΔProx) was similar to that of the ΔGRK5/6 cells (∼70% of the WT response). Together the data suggest that the relative importance of the phosphorylation clusters for β-arrestin binding is distal>proximal>terminal, giving an explanation for why GRK5 appears to phosphorylate ACKR3 more efficiently than GRK2 and dominates β-arrestin recruitment (Fig. 2). Additionally, and as discussed below, GRK5 does not require Gβγ for efficient phosphorylation, whereas GRK2 does^21^. Because ACKR3 does not activate G proteins^9^ the contribution of GRK2 is therefore predictably limited.

We also investigated the effects of the phosphorylation cluster mutations on CXCL12-induced active internalization of ACKR3 by BRET. The ΔDistal mutant receptor showed significant inhibition, whereas ΔTerm and ΔProx were similar to WT ACKR3 (Fig. 4C). The GRK5 specific cluster construct (ΔProx/ΔDistal) showed even further impairment of the internalization (less than 10% change in surface receptor, Fig. 4D) while the triple cluster knockout ΔPDT mutant showed no agonist-induced active internalization. By contrast, the GRK2-like cluster mutant, ΔDistal/ΔTerm (Fig. 4D) showed similar internalization as the ΔDistal construct (Fig. 4C). These results suggest that, as for arrestin recruitment, the distal phosphorylation sites are most important for efficient active internalization. Mutations of the proximal and terminal sites only showed a significant effect when combined with the distal motif, with the proximal being more impactful than the terminal sites.

In contrast to agonist-mediated active ACKR3 internalization, scavenging of CXCL12 was not significantly affected by mutation of the individual phosphorylation clusters, including the ΔDistal mutation (Fig. 4E). Only the ΔPDT mutant and the ΔProx/ΔDistal mutant showed significantly reduced chemokine uptake. The amount of CXCL12 removed by the ΔPDT and ΔProx/ΔDistal mutants was similar to the chemokine uptake by WT ACKR3 in ΔGRK2/3/5/6 and ΔGRK5/6 cells, respectively, again indicating the dominance of GRK5.

Finally, we also examined the effects of the phosphorylation mutations on constitutive (passive) internalization. As with the ΔGRK2/3/5/6 cells, the triple cluster phosphorylation mutation construct (ΔPDT) had no effect (Fig. 4F). The independence of constitutive internalization and dependence of scavenging on ACKR3 phosphorylation is consistent with a previous study^11^. Altogether our data suggests that chemokine scavenging by ACKR3 is mediated by both a phosphorylation-dependent internalization process (dominated by GRK5 in HEK293A cells) and to a lesser extent by a phosphorylation-independent constitutive internalization process. The phosphorylation dependence led us to consider the role of β-arrestin.

### β-arrestins are dispensable for ACKR3 internalization and scavenging

Phosphorylation of GPCRs triggers their interaction with β-arrestins, which then often facilitate receptor internalization. Because internalization is a key requirement of scavenging, and ACKR3 scavenging is dependent on phosphorylation, a logical hypothesis would be that β-arrestins mediate the process. However, the role of β-arrestins in scavenging has been controversial, with some reports suggesting arrestins are essential for ACKR3 internalization and scavenging^17, 25^ and others suggesting they are dispensable^11, 16, 18^. To assess the role of β-arrestin in our system, we conducted experiments in β-arrestin1/2-knockout (ΔArrb) HEK293 cells^26^. As shown in Fig. 5, CXCL12-mediated active internalization (Fig. 5A) and CXCL12 scavenging (Fig. 5B) show little difference in ΔArrb cells compared to WT cells. Likewise, constitutive (passive) internalization was not impacted by the loss of arrestins (Fig. 5C). Thus, although β-arrestin is recruited to the receptor, it has only a minor effect on agonist-induced internalization and chemokine uptake, suggesting a role in an as yet to be identified function.

**Figure 5:**
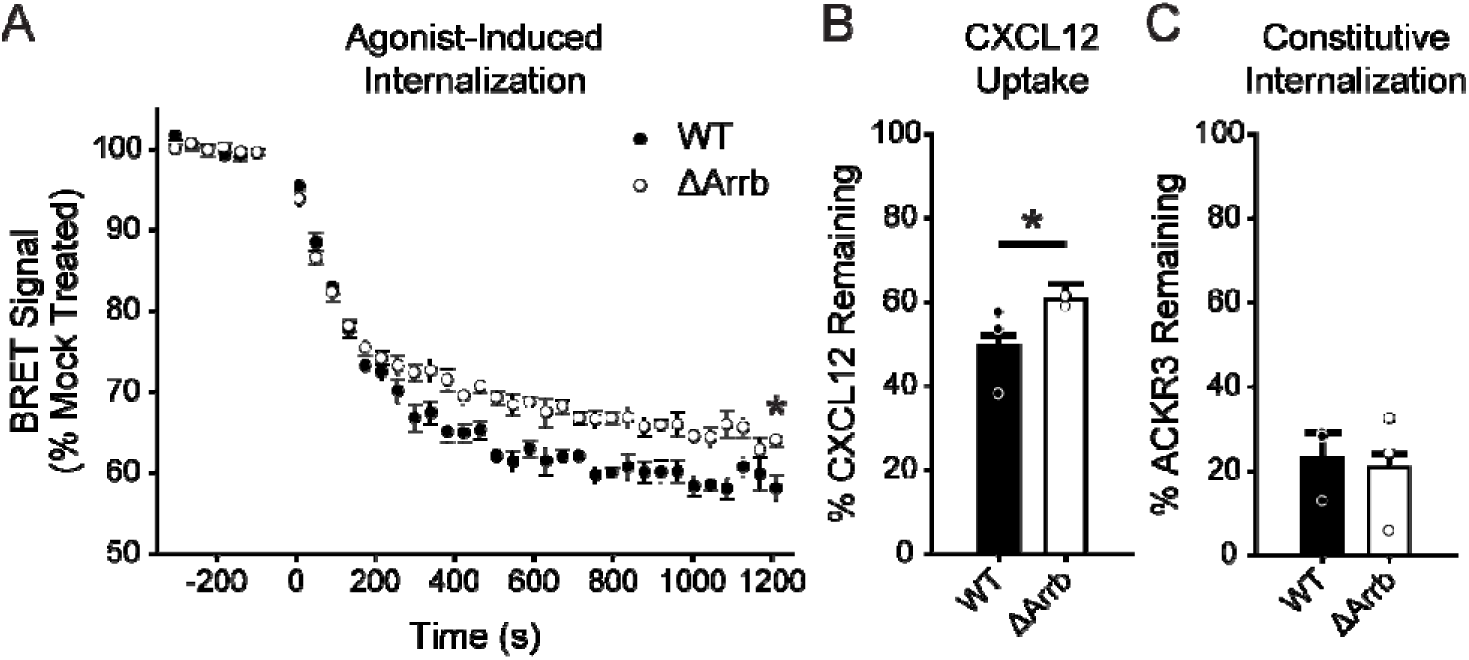
β-arrestins are dispensable for ACKR3 internalization and CXCL12 scavenging in HEK293 cells. A) CXCL12-mediated, active internalization in WT and ΔArrb HEK293 cells measured by decreasing BRET between rGFP_CAAX and ACKR3_RlucII. Note that the cellular background for ΔArrb is HEK293 which leads to slightly different responses compared to the HEK293A cells used in all other figures. B) Chemokine uptake by WT ACKR3 in ΔArrb and WT HEK293 cells. Remaining chemokine was quantified by ELISA and compared to non-ACKR3 expressing cells. C) ACKR3 agonist-independent, passive internalization in WT and ΔArrb HEK293 cells. Data presented are the composite of three independent experiments measured in triplicate and errors represent standard deviations. Statistical significance was determined by one-way ANOVA followed by Bonferroni test. *P<0.05.

### G**βγ** enhances GRK2-mediated responses of ACKR3

Although GRK5 was the dominant kinase in our HEK293A system, other reports suggest that GRK2 plays an important role in driving ACKR3 responses in HEK293 cells and primary neurons^11, 16^. One explanation for the observed dominance of GRK5 in our experiments may be the inability of ACKR3 to activate G proteins and release Gβγ^9, 27, 28^, because Gβγ is required for GRK2 membrane recruitment and efficient receptor phosphorylation^21, 29, 30^. In contrast, GRK5 is constitutively associated with membrane phospholipids and does not bind or require Gβγ^31, 32^. To test the hypothesis that GRK2 phosphorylation of ACKR3 is enhanced by increasing free Gβγ, we utilized β-arrestin recruitment to ACKR3 as an indirect measure of the phosphorylation status of the receptor. In WT cells co-transfected with Gβ^1^γ^2^ along with ACKR3_RlucII and βarr2_GFP, β-arrestin recruitment was enhanced by approximately 50% (area under the curve) compared to cells without additional Gβγ (Fig. 6A,B). The effect of Gβγ addition in ΔGRK5/6 cells was even greater, increasing from ∼36% of WT without extra Gβγ to 84% of WT cells with the co-transfection (Fig. 6B,C). Over-expression of the C-terminal domain of GRK3 (GRK3-CT), which inhibits GRK2/3 phosphorylation by sequestration of free Gβγ^30, 33, 34^, suppressed β-arrestin recruitment in WT cells (Fig. 6A) and nearly eliminated the interaction in ΔGRK5/6 cells (Fig. 6C). Neither over-expression of Gβγ nor GRK3-CT had an effect on β-arrestin recruitment in ΔGRK2/3 cells, indicating that these treatments were specific to GRK2 and not GRK5 (Fig. 6B).

**Figure 6:**
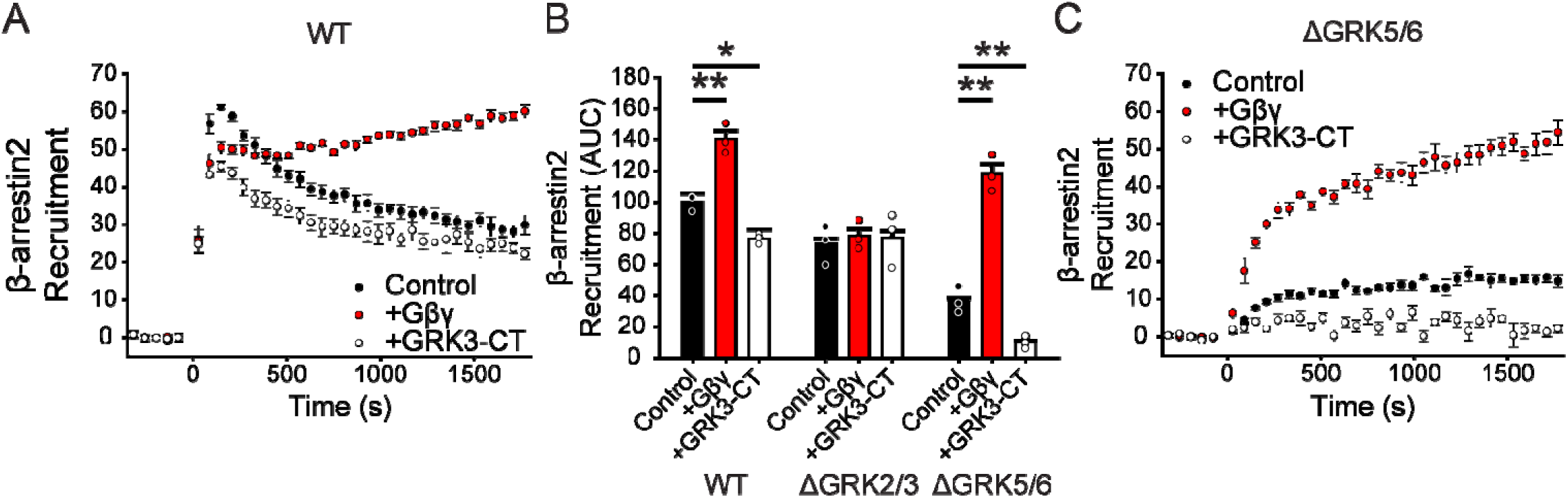
GRK2 phosphorylation of ACKR3 is enhanced by expression of Gβγ in HEK293A cells. A) GFP_βarr2 recruitment to ACKR3_RlucII measured by BRET after 100 nM CXCL12 addition at 0 min in WT HEK293A cells co-transfected with Gβγ or GRK3-CT. B) Quantification of Barr2 recruitment in WT, ΔGRK2/3 and ΔGRK5/6 cells with co-transfection with Gβγ or GRK3-CT by integration of the area under the BRET curves after chemokine addition. Areas are normalized to WT ACKR3 recruitment in WT cells without additional treatment. C) Recruitment of β-arrestin2 to ACKR3 in ΔGRK5/6 cells with Gβγ and GRK3-CT co-expression. Data represent three independent experiments measured in triplicate and individual experiments are presented as points in (B). Errors are reported as standard deviation and statistical significance was determined by one-way ANOVA followed by a Bonferroni test. *P<0.01, **P<0.001.

We next tested if Gβγ enhanced GRK2 activity also improves ACKR3 active internalization and chemokine scavenging. In WT cells, Gβγ co-expression had no effect on agonist-induced active ACKR3 internalization (Fig. 7A), however, a modest increase in internalization was observed in ΔGRK5/6 cells with added Gβγ (Fig. 7B). Likewise, CXCL12 scavenging was significantly enhanced in ΔGRK5/6 cells with added Gβγ, but not in WT cells (Fig. 7C). Together these results are consistent with efficient active internalization and scavenging by ACKR3 phosphorylated by GRK5. However, in the absence of GRK5, GRK2 plays a more important role and requires free Gβγ for efficient activity. Finally, to ascertain that the observed dominance of GRK5 was not a consequence of insufficient GRK2, we performed the BRET-based, β-arrestin2 recruitment experiment with CCR2, a CCKR which is known to be phosphorylated by GRK2/3^35^. CCL2-activated CCR2 showed robust association with β-arrestin2 in WT and ΔGRK5/6 cells indicating ample phosphorylation by GRK2/3 (Fig. 8A,B). In fact, the recruitment profiles to CCR2 were similar to those of ACKR3 in ΔGRK5/6 cells when Gβγ was present, including a prolonged association of the receptors with β-arrestin2 (Fig. 6, Fig. 8). In ΔGRK2/3 cells, β-arrestin2 recruitment to the two receptors was also nearly identical (Fig. 8C). Together these data suggest that the observed dominance of GRK5 in ACKR3 phosphorylation is largely due to a lack of free Gβγ and not a limitation of GRK2 expression or kinase preference for ACKR3.

**Figure 7:**
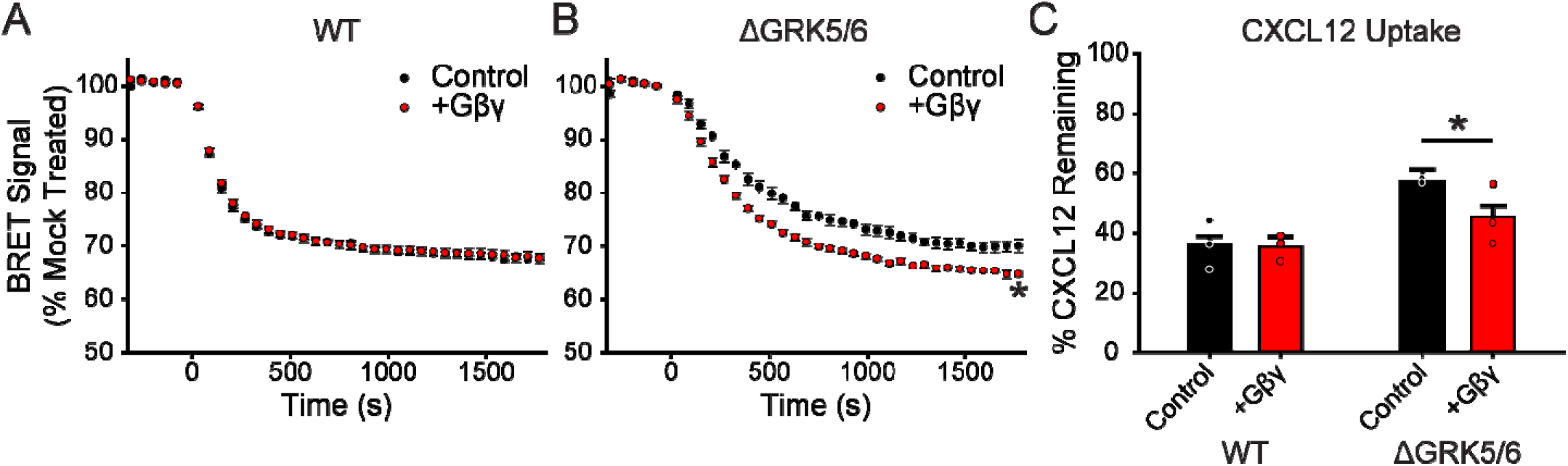
Gβγ co-transfection enhances ACKR3 internalization and CXCL12 scavenging, but only in the absence of GRK5/6 in HEK293A cells. A,B) ACKR3_RlucII active internalization was tracked by rGFP_CAAX BRET with and without co-transfection of Gβγ subunits in WT (A) and ΔGRK5/6 (B) cells following addition of 100 nM CXCL12. Plots are the average of three independent experiments measured in triplicate. C) CXCL12 uptake in WT and ΔGRK5/6 cells with Gβγ co-transfection. Data from cells without extra Gβγ are repeated here from Fig. 3 for comparison. Errors were reported as standard deviations and statistical significance was determined by one-way ANOVA followed by a Bonferroni test. *P<0.001.

**Figure 8:**
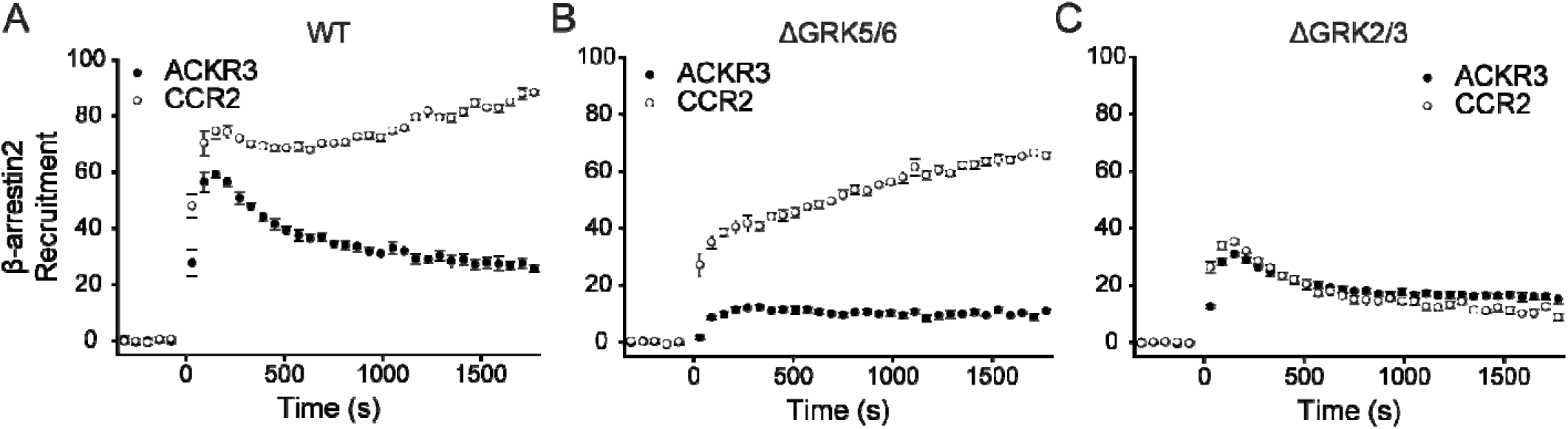
β-arrestin2 recruitment to the canonical GPCR CCR2 is mediated primarily by GRK2/3. A-C) GFP_βarr2 recruitment to ACKR3_RlucII or CCR2_rlucII after stimulation with either 100 nM CXCL12 or CCL2, respectively, was tracked by BRET in WT (A), ΔGRK5/6 (B), and ΔGRK2/3 (C) cells. Data is the average of three independent experiments performed in triplicate and errors reflect standard deviations.

### Co-activation of CXCR4 enhances GRK2-mediated responses of ACKR3

Because ACKR3 does not activate G proteins, we hypothesized that there must be another source of Gβγ in order for GRK2 to contribute to ACKR3 phosphorylation in WT and ΔGRK5/6 cells. Low endogenous levels of CXCR4 have been observed in HEK293 cells^36^. Accordingly, we first tested if the small molecule CXCR4 antagonist, IT1t^37^, would suppress GRK2 phosphorylation and β-arrestin recruitment to ACKR3 by virtue of blocking Gβγ release by CXCL12-activated CXCR4. As expected, IT1t had no effect in the ΔGRK2/3 cells. However, there was a significant decrease in β-arrestin recruitment to ACKR3 with IT1t treatment in WT and ΔGRK5/6 cells (Fig. 9A,B), consistent with the presence and contribution of endogenous CXCR4. As a side note, ACKR3 has been widely reported to signal through β-arrestin, resulting in phosphorylation of ERK1/2 and AKT^9, 10, 38^, although more recent studies have called these observations into question^7^. As shown in Supplementary Fig. 3, we observed that CXCL12-mediated phosphorylation of ERK is inhibited by IT1t, suggesting that it is also due to a small level of endogenous CXCR4 and not ACKR3.

**Figure 9:**
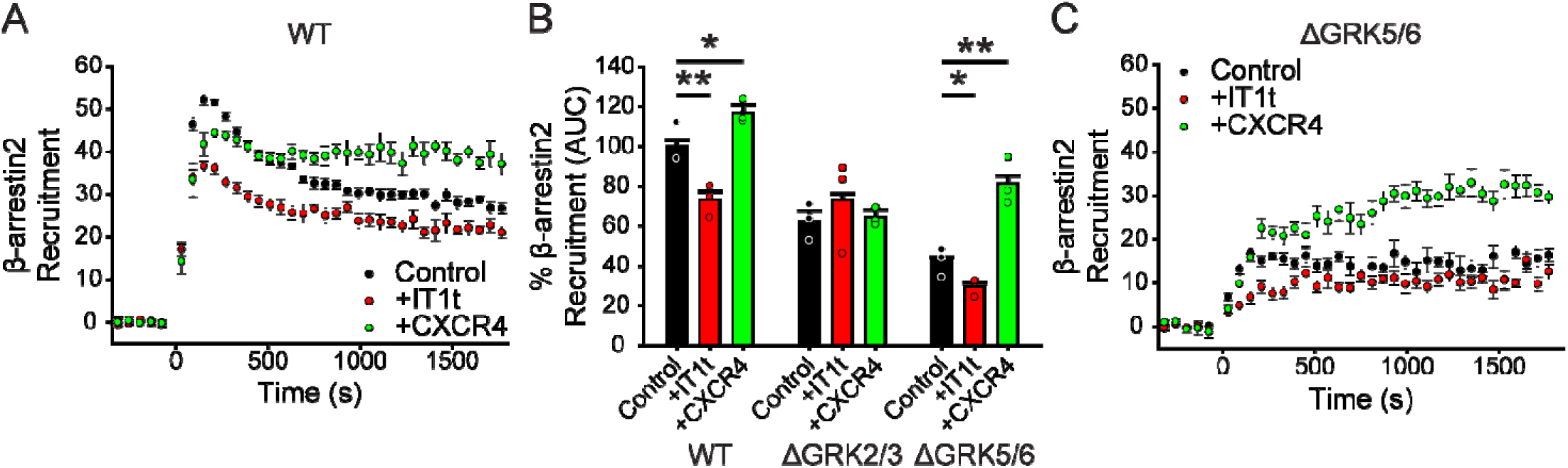
Concurrent CXCR4 activation increases ACKR3 phosphorylation by GRK2/3. A) GFP_βarr2 recruitment to ACKR3_RlucII after 100 nM CXCL12 addition in WT cells treated with 100 µM IT1t for 45 min before the experiment or co-transfected with additional CXCR4 DNA. Individual points are the average of three experiments measured in triplicate. B) Area under the curve analysis of GFP_βarr2 recruitment time courses after CXCL12 addition normalized to WT ACKR3_RlucII in WT cells without treatment. C) Recruitment time courses of GFP_βarr2 to ACKR3_RlucII by BRET in ΔGRK5/6 cells treated identically to the WT cells described in (A). Errors were represented by standard deviation and statistical significance determined by one-way ANOVA followed by a Bonferroni test. *P<0.005, **P<0.001.

Finally, we also tested if exogenous expression of CXCR4 could enhance GRK2-mediated ACKR3 activities. As shown in Fig. 9A and B, addition of CXCR4 slightly increased CXCL12-mediated β-arrestin2 recruitment to ACKR3 in WT cells. However, in ΔGRK5/6 cells, exogenous CXCR4 doubled the recruitment BRET, presumably by liberating Gβγ and enhancing GRK2 activity (Fig. 9B,C). The ability of CXCR4 activation to regulate the activity of ACKR3 may be the source of some of the discrepancies related to ACKR3 function reported in the literature^9, 14, 15, 18, 39^. It also provides a means of crosstalk between the two receptors.

## Discussion

Both GRK2 and GRK5 have been reported to phosphorylate the C-terminus of ACKR3^11, 25^, a post translational modification that is counterbalanced by ubiquitination in regulating ACKR3 levels and scavenging capacity^19, 20^. However, only GRK2 has been reported to promote β-arrestin interaction, receptor internalization, and chemokine uptake^11, 16^, whereas the relative contribution of GRK5 has not been systematically studied. Here we undertook a comparative study of these two ubiquitously expressed kinases to understand their installed phosphorylation patterns, their relative efficiencies, and their role in ACKR3 activities. We were also interested in understanding how GRK2 regulates ACKR3, given that GRK2 depends on Gβγ for efficient phosphorylation, yet most studies suggest ACKR3 does not activate heterotrimeric G proteins, necessary for Gβγ release. Because GRK5 functions in a G protein independent manner, it is expected to more efficiently phosphorylate ACKR3 than GRK2, all else being equal. This turned out to be the case in our studies using HEK293A cells where GRK5 dominated ACKR3 phosphorylation over GRK2. Phosphorylation by GRK2 was more limited and observed most strongly as a consequence of CXCR4 and ACKR3 co-stimulation by CXCL12. Additionally, supplementation of CXCR4 increased phosphorylation of ACKR3 by GRK2 whereas IT1t antagonism of CXCR4 suppressed it. These results suggest a means for ACKR3 to ‘sense’ CXCR4 activation and respond by increasing chemokine uptake (Fig. 10). By including a role for CXCR4 in the phosphorylation process, this mechanism expands the competing phosphorylation-ubiquitination feedback regulation of ACKR3 by extracellular CXCL12 levels proposed by Wong et al^19^. Excess CXCL12 activates additional CXCR4, which in turn promotes chemokine scavenging by ACKR3. ACKR3/CXCR4 crosstalk would obviously require that the receptors are expressed in the same cell, as observed in cortical interneurons^11^. Alternatively, in cells with low CXCR4 expression, such as in our HEK293A cells, ACKR3 phosphorylation requires a Gβγ independent kinase like GRK5. The results also suggest that the phosphorylation of ACKR3 by GRK2 reported in earlier studies^11, 16^ is likely promoted by endogenous CXCR4.

**Figure 10:**
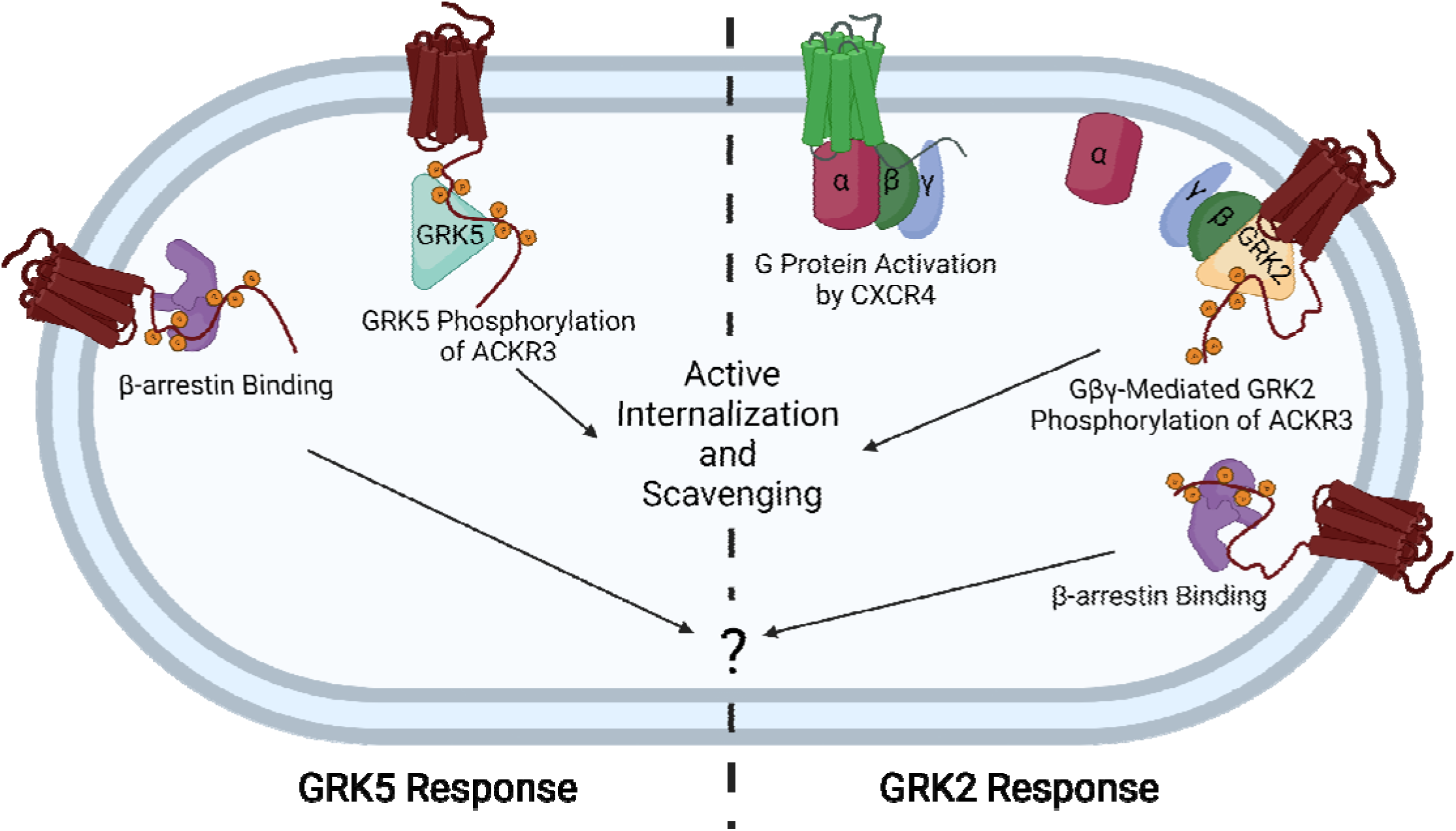
ACKR3 GRK specificity is dependent on whether CXCR4 is co-activated or not. When expressed alone, the atypical receptor is primarily phosphorylated by GRK5, which drives agonist-mediated internalization and scavenging. In the context of CXCR4 co-expression, the co-activation of CXCR4 activates heterotrimeric G proteins and releases Gβγ, which recruits GRK2 and mediates phosphorylation of both receptors. This allows for ACKR3 to sense the activation of CXCR4 and then enhance the scavenging, decoy responses of ACKR3. This image does not include the passive, constitutive internalization and scavenging pathway. Image created with BioRender.

An obvious question is whether GRK discrimination extends to other receptor systems that are biased away from G proteins. Consistent with the bias model, Shukla and colleagues demonstrated that the atypical receptors C5a receptor subtype 2 (C5aR2) and ACKR2 both show a decrease in β-arrestin recruitment in ΔGRK5/6 cells, whereas ΔGRK2/3 cells have little to no impact^23^. In contrast, canonical, G protein-activating receptors like CCR2 (Fig. 8)^35, 40^ and µ-opioid receptor^41^ are both preferentially phosphorylated by GRK2/3. The behavior also appears to extend to biased ligands. When angiotensin II (AngII) type-1 receptor (AT1R) is stimulated by the neutral agonist AngII, phosphorylation is mediated by GRK2/3. However, the arrestin-biased ligand TRV027 completely shifts this dependence to GRK5/6, consistent with G protein activation mediating GRK discrimination^42^. In an exception to this paradigm, ACKR4, which does not couple G proteins, exclusively associates with Gβγ-regulated GRK2 and GRK3^43^. This suggests that a novel interaction may promote GRK2/3 phosphorylation that is distinct from the canonical mechanism or that another CCKR responds to the ACKR4 ligands CCL19, CCL21, and CCL25 and supplies Gβγ. Although it is tempting to propose this GRK regulation model as general feature of arrestin-biased systems, more detailed studies are needed to fully understand how extensively it applies and to explain exceptions such as ACKR4.

The availability of GRKs has been proposed as a mechanism for modulating GPCR function in different cellular systems^44^. Functional differences due to GRK phosphorylation has been demonstrated for many receptors including CXCR4^45^, β2 adrenergic receptor^46^, and others^23, 41, 42, 47^. Our results suggest a unique twist where the different GRKs alter the regulation of a given function, in this case chemokine scavenging by ACKR3, with potential consequences on the activity of another receptor. Specifically, when the receptor is expressed alone, GRK5 drives chemokine-mediated phosphorylation of ACKR3. However, in the presence of CXCR4, phosphorylation by GRK2 is enhanced, which in turn may limit the availability of CXCL12 to CXCR4, leading to effects on CXCR4 signaling, desensitization, and degradation^11, 48^. Whether phosphorylation by GRK2 vs. GRK5 also leads to differences in ACKR3 functional responses remains to be determined. However, it is notable that GRK2 mediated phosphorylation leads to different phosphorylation patterns on the ACKR3 C-terminus and more persistent complexes with β-arrestin than GRK5 (see Fig. 4 and 6).

Despite the requirement of phosphorylation for agonist-induced internalization of ACKR3, scavenging was not completely abolished in the ΔGRK2/3/5/6 cells and when all C-terminal phosphorylation sites in ACKR3 were removed. Rather, low level (20-30%) chemokine uptake persisted in both cases. Constitutive internalization and recycling of ACKR3 was also fully retained under these conditions. These data suggest that in addition to the active, agonist-induced phosphorylation-dependent pathway, an alternative passive, constitutive phosphorylation-independent mechanism exists for transporting chemokine into cells, a mechanism that is not unprecedented. For example, we recently demonstrated that scavenging by CCR2, a “dual function” chemokine receptor that also couples to G proteins, continues in the absence of phosphorylation^40^. Furthermore, the low level uptake of chemokine by phospho-deficient ACKR3 may explain observations that, whereas ACKR3 knockout mice are embryonically lethal^49^, mice expressing a phosphorylation-deficient ACKR3 mutant are viable^11^. Nevertheless, the phospho-deficient ACKR3 mice showed defects in neuronal CXCR4 expression and responsiveness to CXCL12 as well as cell migration, suggesting that constitutive scavenging is unable to fully regulate chemokine levels^11^. Along these lines, zebrafish primordium tissue expressing phospho-deficient ACKR3 still migrated normally when exposed to physiological CXCL12 levels, but migration was compromised with elevated “chemokine floods”^19^. Together, these data suggest that constitutive internalization and recycling of ACKR3 may effectively regulate low levels of chemokine even in the absence of phosphorylation, but receptor phosphorylation is required to accommodate chemokine surges. Indeed, Wong et al. showed support for this hypothesis by demonstrating that phosphorylation increases ACKR3 levels and scavenging efficiency by enhancing plasma membrane recycling rather than degradation^19^.

Due to the β-arrestin bias of ACKR3, coupled with the role of β-arrestin in regulating the internalization of many GPCRs, the expectation was that scavenging by ACKR3 would occur in a β-arrestin dependent manner. However, the role of β-arrestin in scavenging has been controversial with some reports involving HEK293 and mouse embryonic fibroblasts indicating it is necessary for ACKR3 internalization and chemokine scavenging^17, 25^, and others involving mouse neurons and HEK293 cells showing no impact^11, 16, 18^. Our data also suggests that constitutive and agonist-induced internalization, as well as scavenging of CXCL12, occur largely independent of β-arrestin. ACKR3 has been shown to signal through ERK/MapK/AKT pathways via β-arrestins^9, 10, 38, 50^, although this appears to be limited to cells with currently active heterotrimeric G protein signaling^51^. Likewise, we demonstrate that ERK/MapK phosphorylation in our HEK293 cells is CXCR4 dependent (Supplementary Fig. 3). Because ACKR3 readily recruits β-arrestin in response to almost every ligand^27^, it likely plays a critical role in some common ACKR3-mediated process, but that role remains a mystery.

In conclusion, our study reveals differences in ACKR3 regulation in response to the activity of GRK2 and GRK5. GRK5 dominates when ACKR3 binds CXCL12 in isolation, whereas GRK2 phosphorylates ACKR3 when CXCR4 and ACKR3 are co-stimulated by CXCL12, providing a readout of CXCR4 activation status, increasing CXCL12 scavenging, and possibly modulating CXCR4 responses and trafficking. Because ACKR3 is also a receptor for the chemokine CXCL11, a similar crosstalk may exist in cells co-expressing CXCR3. Crosstalk with other receptors that do not share a common ligand is also conceivable.

## Materials and Methods

### Materials

Unless otherwise stated all chemicals and reagents were purchased from Sigma and Fisher. HEK293A WT, HEK293 WT, and CRISPR GRK and β tin1/2-knockout cells were generous gifts from Aska Inoue (Tohoku Univ.)^23, 42^. Methoxy e-Coelenterazine (Prolume Purple) was purchased from Nanolight Technologies (Prolume LTD) and fluorescent antibodies from Li-COR Biosciences. CXCL12 ELISA kits were from R&D Systems.

### Cloning

N-terminally FLAG-tagged human ACKR3 (residues 2-362) was cloned into pcDNA3.1 expression vector either alone (FLAG_ACKR3) or followed by a C-terminal fusion of *Renilla* luciferase II (ACKR3_RlucII) in the pcDNA3.1 vector for use in BRET assays. For purified receptor studies, ACKR3 (residues 2-362) with an N-terminal HA signal sequence was cloned into pFasBac vector followed by C-terminal 10His and FLAG tags. Site-directed mutagenesis was performed by overlap extension and confirmed by Sanger sequencing.

### CXCL12 Purification from *E. Coli*

The chemokine CXCL12 was expressed and purified as previously described^52^. Briefly, mature chemokine sequence, preceded by an 8His tag and an enterokinase cleavage site, was cloned into a pET21-based vector and expressed in BL21(DE3)pLysS cells by IPTG induction. Cells were lysed by sonication and chemokine containing inclusion bodies were dissolved in 50 mM Tris, 6 M guanidine-HCl, 50 mM NaCl, pH 8.0. The chemokines were bound to a Ni-NTA column, washed with 50 mM MES, 6 M guanidine-HCl, 50 mM NaCl, pH 6, and eluted with 50 mM acetate, 6 M guanidine-HCl, 50 mM NaCl, pH 4. CXCL12 was refolded in 50 mM Tris, 500 mM arginine-HCl, 1 mM EDTA, 1 mM glutathione disulfide, pH 7.5 before removal of the tag by enterokinase. The cleaved material was then bound to a C18 HPLC column (Vydac) (buffer A: 0.1% trifluoroacetic acid; buffer B: 0.1% trifluoroacetic acid, 90% acetonitrile) and eluted by a linear gradient of buffer B from 33-45%. The peak was collected, lyophilized, and stored at -80°C until use.

### Arrestin Expression and Purification

Expression and purification of β-arrestin1/2 was described previously^53^. Briefly, the pTrcHisB plasmid containing bovine β-arrestin1 and β-arrestin2 (1-393) were transformed into *E. coli* Rosetta cells and protein expression was induced with the addition of 25 μM (β-arrestin1) and 35 μM IPTG (β-arrestin2) for 4 hours at 30 °C. Cells were collected by centrifugation and the pellet was processed immediately or stored at -80 °C. The cell pellets were resuspended and homogenized in 20 mM MOPS pH 7.5, 200 mM NaCl, 5 mM EDTA, 2 mM DTT, 1 mM PMSF, and leupeptin, lima bean trypsin protease inhibitor. Cells were lysed using an Avestin C3 emulsifier. The lysate was clarified by centrifugation at 18,000xg for 60 min. The supernatant was collected and arrestin was precipitated by the addition of (NH_4_)_2_SO_4_ to a final concentration 0.32 mg/ml. Precipitated arrestin was collected by centrifugation at 18,000xg for 90 min and dissolved in buffer containing 20 mM MOPS (pH 7.5), 2 mM EDTA, and 1 mM DTT, then centrifuged again at 18,000 for 60 min to remove insoluble parts. The supernatant containing soluble arrestin was applied onto a heparin column and eluted with a linear NaCl gradient (0.2-1 M). Fractions containing arrestin were identified by SDS-PAGE and combined. For β-arrestin1, the salt concentration of the pooled fractions were adjusted to 50 mM, loaded onto a 5 ml HiTrap Q HP column (Cytiva), and eluted with a linear NaCl gradient. For β-arrestin2, the salt concentration of the pooled fractions was adjusted to 100 mM, and the solution was loaded onto a linked 1 ml HiTrap Q HP and 1ml HiTrap SP HP column. β-arrestin2 flows through the Q column but binds the SP column. The columns were uncoupled and a linear NaCl gradient was used to elute β-arrestin2 from the SP column. The fractions containing arrestin were identified by SDS-PAGE and combined, concentrated with a 30 kDa cutoff Amico concentrator to ∼500 μl, then further purified using a Superdex 200 increase 10/300 GL column (Cytiva) equilibrated with 20 mM MOPS (pH 7.5), 150 mM NaCl, and 0.5 mM TCEP. The peak fractions were collected, concentrated with a 30 kDa cutoff Amicon concentrator, and stored at -80 °C.

### GRK Purification

GRK2 and GRK5 expression and purification were described previously^54, 55^. Briefly, a pMAL plasmid containing human full-length GRK5 with C terminal 6his tag were transformed into *E. coli* Rosetta cells. The expression of GRK5 was induced by the addition of 200 μM IPTG at OD around 0.6-0.8 and the cultures were incubated with shaking at 18 °C overnight. For purification cell pellets were resuspended and homogenized in lysis buffer (20 mM HEPES pH 8.0, 400 mM NaCl, 0.1% Triton-X (v/v), 2 mM DTT, DNase, 0.1 mM PMSF and leupeptin, lima bean trypsin protease inhibitor). The cells were then lysed using an Avestin C3 emulsifier and centrifuged at 18,000 rpm for 30 min. The supernatant was combined and loaded onto a 3 ml home-packed Ni^2+^-NTA column pre-equilibrated with buffer A (20 mM HEPES pH 8.0, 400 mM NaCl and 0.5 mM DTT). The column was then washed with 50 ml buffer A, followed by 100 ml buffer B (20 mM HEPES pH 8.0, 100 mM NaCl and 0.5 mM DTT plus 20 mM imidazole). The bound protein was eluted in approximately 2 ml fractions with buffer B plus 200 mM imidazole. The purity of GRK5-HIS after this step was about 60% as revealed by Coomassie blue staining of samples assessed via SDS–PAGE. The fractions were pooled and loaded onto a linked 1 ml HiTrap Q HP and 1 ml HiTrap SP HP column. GRK5-HIS flows through Q column and binds to the SP column. The columns were uncoupled and a linear NaCl gradient (0.1-0.6 M) was used to elute GRK5-HIS from the SP column. GRK5 elutes with ∼0.3-0.5 M NaCl. The fractions containing GRK5-HIS were identified by SDS-PAGE and combined, concentrated with a 50 kDa cutoff Amicon concentrator to ∼500 μl, then further purified using a Superdex 200 increase 10/300 GL column equilibrated with 20 mM HEPES (pH 8.0), 100 mM NaCl, and 0.5 mM TCEP. The peak fractions were collected, concentrated with a 50 kDa cutoff Amicon concentrator, and stored at - 80°C.

Human GRK2 S670A with a C-terminal hexahistidine tag was expressed in *Spodoptera frugiperda* (*Sf*9) cells using the Bac-to-Bac insect cell expression system (Life Technologies). The insect cells were harvested 48 hours post-infection and homogenized with buffer containing 20 mM HEPES pH 8.0, 400 mM NaCl, 2 mM DTT, 1 mM PMSF, and leupeptin, lima bean trypsin protease inhibitor. Cells were lysed using an Avestin C3 emulsifier. The lysate was clarified by centrifugation at 18,000xg for 60 min. GRK2 was purified from the clarified lysate as described above for GRK5 using nickel-nitrilotriacetic acid affinity chromatography. The purity of GRK2-HIS after this step was ∼90% as revealed by Coomassie blue staining of samples assessed via SDS–PAGE. Fractions containing GRK2 were pooled and further purified on a Superdex 200 increase 10/300 GL column equilibrated with 20 mM HEPES (pH 8.0), 100 mM NaCl, and 0.5 mM TCEP. The peak fractions were collected, concentrated with a 50 kDa cutoff Amicon concentrator, and stored at -80°C.

### ACKR3 Expression and Purification from *Sf*9 Cells

Expression and purification of ACKR3 from *Sf*9 cells was performed as previously described^27, 56^. Briefly, Sf9 cells were transfected with ACKR3 or CXCL12_LRHQ_, a high-affinity variant of CXCL12 with residues 1-3 replaced by the motif LRHQ, in pFasbBac vectors using X-tremeGene transfection reagent (Roche) to produce baculovirus. The receptors and CXCL12_LRHQ_ were co-expressed by infecting Sf9 cells at a density 2x10^6^ cells/ml with a multiplicity of infection (MOI) of 6 for each virus.

After 48 hours, cell pellets were collected and stored at -80 °C. Membranes were prepared by dounce-homogenization in hypotonic buffer (10 mM HEPES pH 7.5, 10 mM MgCl_2_, 20 mM KCl) and then three times with hypotonic buffer with 1 M NaCl. Between each cycle of douncing, the membranes were pelleted by centrifugation at 50k x g for 30 min and resuspended. The prepared membranes were then solubilized in 50 mM HEPES pH 7.5, 400 mM NaCl, 0.75%/0.15% dodecyl maltoside/cholesteryl hemisuccinate (DDM/CHS) with 2 mg/ml iodoacetamide and a Protease Inhibitor tablet (Roche). After 4 hours, the insoluble material was removed by 50k x g centrifugation for 30 min and the supernatant was added to Talon resin (Clontech) with 20 mM imidazole to bind overnight at 4°C. The resin was transferred to a column and washed with WB1 (50 mM HEPES pH 7.5, 400 mM NaCl, 0.1/0.02% lauryl maltose neopentyl glycol (LMNG)/CHS, 10% glycerol, 20 mM imidizole), followed by WB2 (WB1 with 0.025/0.005% LMNG/CHS), and finally eluted with EB (WB2 with 250 mM imidizole). The elutions were pooled and concentrated to 500 µl before passing over a PD MiniTrap G-25 desalting column (GE Healthcare) equilibrated with 50 mM HEPES pH 7.5, 100 mM NaCl, 0.025/0.005% LMNG/CHS, 10% glycerol. The final protein concentration was calculated using an A_280_ extinction coefficient of 85000, snap frozen in liquid nitrogen, and stored at -80°C until use.

### *In Vitro* Pulldown of Arrestin by ACKR3

To phosphorylate the receptors, 1.1 µM of purified ACKR3:LRHQ complex was incubated with 1.1 µM CXCL12_LRHQ_ and either 1.1 µM of GRK2 or GRK5 in 50 mM HEPES pH 8.0, 10 mM MgCl_2_, 0.1% LMNG, 0.005% 1,2-dioctanoyl-sn-glycero-3-phospho-1’-myo-inositol-4’,5’-bisphosphate (C8-PIP2) (Avanti) with 1 mM ATP for 20 min at room temperature. Next, purified β-arrestin1 or 2 was added to a final concentration of 2.2 µM and was allowed to complex for 40 min at room temperature. 5 µl of M2 anti-FLAG-resin (Sigma) was then added to the reaction and incubated at 4 °C for 1 hour. The bound complexes were washed in batch 3x with 50 mM HEPES pH 7.5 150 mM NaCl, 0.01/0.001% LMNG/CHS and eluted by adding 250 µg/ml 3xFLAG peptide (final concentration, Sigma). The total supernatant was analyzed via 10% SDS-PAGE. Band densities were quantified using ImageJ software and the amount of arrestin pulled down was reported as a percentage of the amount of ACKR3 in the same experiment after normalizing by molecular weight.

### *In Vitro* ACKR3 Mass Spectrometry

Phosphosite mapping was performed at the Purdue University Proteomics Facility. Briefly, ACKR3 in LMNG or nanodisc was first phosphorylated by GRK2 and GRK5 and then digested with trypsin. The fragments were analyzed via high-resolution MS without TiO_2_ enrichment, and phosphorylation sites identified through peptide ionization patterns compared to the non-phosphorylated primary amino acid sequence.

### Arrestin Binding to ACKR3 Measured by BRET

Recruitment of β-arrestin2 to ACKR3 and CCR2 was measured with a BRET2 assay as previously described^27, 57^. HEK293A cells were initially plated at 750k cells/well in 6-well plates in Dulbecco’s modified eagle media (DMEM) supplemented with 10% fetal bovine serum (FBS) and GlutaMax (Gibco) with 5% CO_2_. The following day, the cells were transfected with 100 ng ACKR3_RlucII DNA (or FLAG_CCR2_rlucII) and 2 µg GFP10_β-arrestin2 (GFP_βarr2, a gift from N. Heveker, Université de Montréal, Canada) both in pcDNA3.1, with 400 ng empty pcDNA3.1 vector using the TransIT-LT1 transfection system (MirusBio) per manufacturer’s protocol and expressed for 40 hours. For experiments with CXCR4, Gβγ, or GRK3-C-terminus (GRK3-CT) co-expressions, transfections were augmented with either 500 ng CXCR4 DNA, 1 µg GRK3-CT (Bovine, residues 547-688; gift from N. Lambert, Augusta Univ.) DNA, or 500 ng each of Gβ_1_ and Gγ_2_ DNA (gift from A. Inoue, Tohoku Univ.) in pcDNA3.1 vector were included and the amount of ACKR3_RlucII and GFP_βarr2 DNA was reduced to 50 ng and 1 µg respectively to accommodate the additional DNA. On the day of the experiment, the cells were washed with PBS (137 mM NaCl, 2.7 mM KCl, 10 mM Na_2_HPO_4_, 1.8 mM KH_2_PO_4_, pH 7.4) and mechanically lifted with Tyrode’s Buffer (25 mM HEPES, 140 mM NaCl, 2.7 mM KCl, 12 mM NaHCO_3_, 5.6 mM Glucose, 0.5 mM MgCl_2_, 0.37 mM NaH_2_PO_4_, pH 7.5) and counted on a Vicell cell counter (Beckman Coulter). 100k cells were plated in each well of a white, clear bottom 96 well plate (BD Falcon) and allowed to re-adhere for 45 min at 37 °C. For IT1t treated samples, the CXCR4 antagonist was added at 100 µM final concentration before re-adhering. Arrestin expression was verified using a Spectramax M5 plate fluorometer (Molecular Devices) with 485 nm excitation, 538 nm emission, and 530 nm cutoff. White backing tape (PerkinElmer) was applied to the plate and the Prolume Purple luciferase substrate was added to a final concentration of 5 µM. Total luminescence was measured using a VictorX Light multilabel plate reader (PerkinElmer) with no filter and an integration time of 0.1 sec. CXCL12 was then added to each well at indicated final concentrations and the plate was read at 410 nm and 515 nm after 20 min incubation at 37 °C. The BRET ratios (515 nm emission/410 nm emission) were baseline matched and normalized to the WT E_max_ measured in the same experiment on the same day. The reported data represents a combined data set of three independent experiments tested in duplicate or triplicate. Points were fitted with a sigmoidal dose-response model using SigmaPlot 11.0 (Systat software Inc.).

The β-arrestin2 binding time courses were set up as described above with the following exceptions. All experiments were read with a TECAN Spark luminometer (Tecan Life Sciences) at 37 °C using default BRET2 settings (blue emission 360-440 nm, red emission 505-575 nm) and 0.5 sec integration time. Experiments were read for 5 min before 100 nM final concentration CXCL12 was added and BRET readings were collected for an additional 30 min. BRET ratios were calculated by red emission/blue emission and the chemokine and mock treated wells were averaged for each experiment. Percent change in BRET due to CXCL12 binding was calculated as the ratio between BRET_CXCL12_ and BRET_Mock_. Data presented represents a combined data set of three independent experiments each performed in triplicate. The area under curves was calculated using GraphPad Prism 9 (GraphPad Software, Inc.).

### ACKR3 internalization measured by BRET

Agonist-mediated internalization of ACKR3 was measured by BRET2 between ACKR3_RlucII and rGFP_CAAX (a gift from M. Bouvier, Université de Montréal) as previously described^24^. Samples were prepared as described for β-arrestin2 recruitment time courses with the exception of the transfected DNA amounts. HEK293 cells were transfected with 42 ng ACKR3_RlucII and 170 ng rGFP_CAAX DNA, with empty pcDNA3.1 to bring the total DNA amount to 2.5 µg/well. Data presented in Figs. 1, 3, and 4C were collected with a PerkinElmer Victor Luminometer, while the time courses in Fig. 4D and 7 were measured on a Tecan Spark luminometer. All settings were identical to those used for arrestin association (described above). Data is presented as percent change compared to mock treated wells and are a composite of three independent experiments. The percent changes after 30 min were compared for statistical analysis.

### Constitutive ACKR3 internalization measured by flow cytometry

ACKR3 was stably expressed in HEK293 cells by lentiviral spinoculation and selection with hygromycin^58^. The stable, homogenous ACKR3 expression was necessary to resolve changes to the surface receptors by flow cytometry. Cells were grown to confluency in 6 cm dishes before washing with cold PBS on ice and lifting with cold accutase (Innovative Cell Technologies, Inc.). Next, 100k cells were transferred to each well of two conical, 96 well plates, one for 37°C experimental samples and the other kept at 4 °C as a control. The cells were washed with cold FACS buffer (PBS, 0.5% bovine serum albumin (BSA)) and labeled with 0.02 µg/well un-conjugated anti-ACKR3 antibody (11G8, R&D Systems) for 1 hour at 4 °C. Unbound antibody was then washed away with FACS buffer. Prewarmed assay buffer (DMEM, 0.5% BSA, 25 mM HEPES pH 7.5) was added to each well of the 37 °C plate and the plate was moved to a 37 °C incubator for 45 min. The 4 °C control plate was left in the 4 °C refrigerator during this step. The samples were then washed with FACS buffer and labeled with 1 µl/well of phycoerythrin-conjugated anti-mouse secondary (F0102B, R&D Systems) 1 hour at 4 °C. Surface ACKR3 was assessed by flow cytometry using a GuavaCyte benchtop flow cytometer (MilliporeSigma)). The geometric mean fluorescence intensity (GMFI) representing the amount of surface labeling for each experiment was quantified using FlowJo software (FlowJo). The constitutive internalization was then represented by the ratio of the GMFI of 37 °C samples to the 4 °C controls measured on the same day.

### ACKR3 scavenging of CXCL12 determined by ELISA

CXCL12 uptake by ACKR3 was quantified by ELISA per manufacturer’s protocols (R&D Systems). Briefly, HEK293 cells were seeded on 6 well plate at 750k cells/well and transfected the next day with 200 ng of ACKR3_RlucII DNA or empty vector per well as above. After 16 hours, the cells were mechanically lifted, counted, and re-plated at 80k cells per well into a 96 well plate and allowed to re-adhere for 6 hours at 37 °C or in a 96-well BRET plate and reattached for 30 min at 37 °C. The BRET plates were washed with Tyrode’s buffer and Prolume Purple was added to a final concentration of 5 µM and the total ACKR3 expression was determined by luminescence. After 6 hours, the media was exchanged for media containing 25 nM CXCL12 and incubated at 37 °C for 16 hours. The media was carefully collected from each well and cellular debris was removed with a 4 min spin at 250 x g. The remaining CXCL12 was detected using an R&D Systems ELISA kit and read using a Spectramax M5 plate reader. The amount of chemokine removed was quantified by the ratio of cells expressing ACKR3 to those transfected with empty vector from the experiments performed on the same day. Three separate experiments were performed in triplicate and averaged together to determine the final amounts of CXCL12 uptake.

### Detection of ERK phosphorylation by western blot

Detection of pERK activation performed as previously described^11^. HEK293 cells were grown to confluency in a 24-well plate and transfected with 200 ng FLAG_ACKR3 per well. After 8 hours, the media was removed and replaced with DMEM without FBS for overnight serum starvation. The wells were treated with 10 nM CXCL12 final concentration at the given timepoints at 37 °C. IT1t treated samples were incubated for 45 min with 100 µM IT1t before beginning the CXCL12 additions. Cells were harvested with hot sample buffer (20 mM Tris pH 6.8, 12.5 mM EDTA, 20% glycerol, 1% SDS, 0.01% bromophenol blue, 100 mM DTT) at 90 °C. Next genomic DNA was sheared by sonication and the samples were boiled at 95 °C for 5 min. The samples were then spun for 1 min at 20k x g and run on a 10% SDS gel before transferring to nitrocellulose membranes (Bio-Rad) and blocking as described above. ERK1/2 phosphorylation was detected by probing with anti-phospho-p44/42 (4370, Cell Signaling) and anti-tubulin (T6074, MilliporeSigma) primary antibodies and detected with fluorescent secondary antibodies (IRDye® 800CW Donkey anti-Mouse and IRDye® 680RD Goat anti-Rabbit, LI-COR Biosciences) using LI-COR fluorescent imaging system. Next the blots were stripped and re-probed with anti-total ERK (06-182, MilliporeSigma) and anti-tubulin and detected using the same secondary as previously. Bands were quantified using ImageJ. The ratio of pERK to total ERK, each adjusted by corresponding tubulin density, was calculated for each lane to correct for differences in loading and staining efficiency. Values were normalized to the 0 min timepoint on the respective membrane before averaging.

### Statistical analyses

Statistical analyses were performed using SigmaPlot 11.0 software and methods described in the figure legends. Bar and symbol representation, along with error bars, are described in figure legends. Unless otherwise noted, scatter plots represent the average of three independent experiments measured in triplicate. For bar charts, the bars represent the average of three independent experiments, whereas the overlaid points report the values from the individual experiments measured in triplicate. All errors are reported as standard deviation. The responses to the CXCL12 titration experiments were fit to sigmoidal dose-response model using SigmaPlot 11.0 and statistical significance was determined using the extra sum-of squares F test. One-way ANOVA follow by Bonferroni t-test was used to determine statistical significance and P values for all other comparisons using SigmaPlot 11.0.

### Data availability

The raw LC-MS data are submitted in MassIVE (<u>massive.ucsd.edu</u>) public database under ID MSV000091173.

## Acknowledgements

We thank M. Bouvier (Université de Montréal), N. Lambert (Augusta Univ.), N. Heveker (Université de Montréal), and Aska Inoue (Tohoku Univ.) for the BRET constructs and cell lines used in our studies. All the mass spectrometry experiments were performed at the Purdue Proteomics Facility, which is administered through the Office of the Executive Vice President for Research. We thank Dr. Uma K. Aryal and other Facility staffs for helping with mass spectrometry sample preparation, data collection and analysis. This work was supported by National Institutes of Health Grants CA254402 (JJGT, TMH), AI161880 (TMH), GM133157 (TMH) CA221289 (JJGT), HL071818 (JJGT), F32 GM137505 (CTS), American Heart Association Post-Doctoral Fellowship 19POST34450193 (QC), and the Walther Cancer Foundation (JJGT).

## Author Contributions

C.T.S. conceived, performed, analyzed most of the experiments and wrote and revised the manuscript. Q.C. performed the *in vitro* phosphorylation for the mass spectrometry experiments, analyzed the mass spectrometry data, and revised the manuscript. J.J.G.T. conceived and supervised the project and revised the manuscript. T.M.H. conceived and supervised the project and wrote and revised the manuscript.

