## Supplemental Materials for "Atypical Chemokine Receptor 3 ‘Senses’ CXC Chemokine Receptor 4 Activation Through GPCR Kinase Phosphorylation"

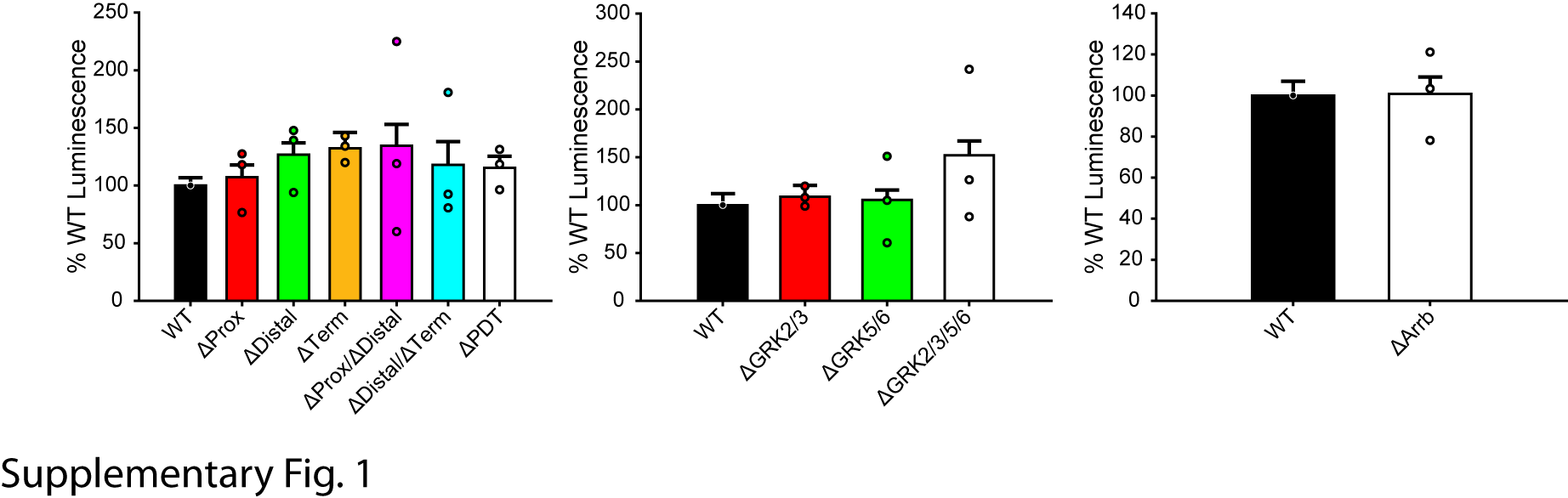

Supplementary Figure 1: Expression of ACKR3_RlucII in HEK293 cells was determined by *Renilla* luciferase luminescence and normalized to WT ACKR3_RlucII expressed in corresponding parental cell lines, HEK293A for phosphorylation deficient ACKR3 and ΔGRK cells and HEK293 for ΔArrb cells. Bars represent average luminescence relative to WT ACKR3 in WT cells and individual experiments presented as points. Errors are reported as standard deviations across three separate experiments in triplicate. None of the tested constructs or cell lines showed significantly different expression level compared to WT ACKR3 in corresponding WT (HEK293A or HEK293) cells.

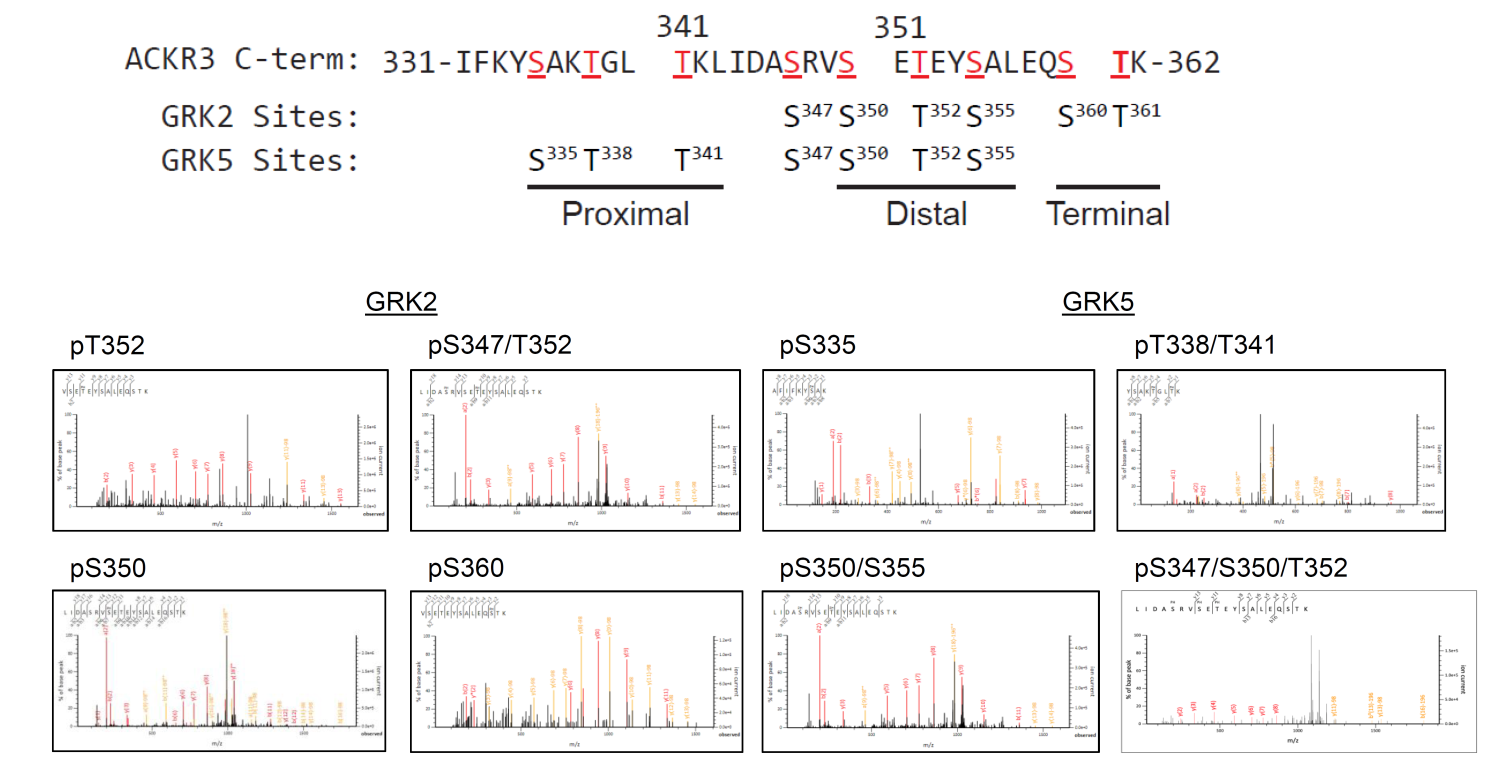

Supplementary Figure 2: GRK2 and GRK5 produce distinct phosphorylation patterns on the C-terminus of ACKR3. The specific phosphate sites identified following phosphorylation by GRK2 or GRK5 are recapitulated from Fig. 4A. Example MS2 spectra of purified ACKR3 in detergent micelles phosphorylated *in vitro* individually by either GRK2 or GRK5 reveal the phosphate positions incorporated by each kinase.

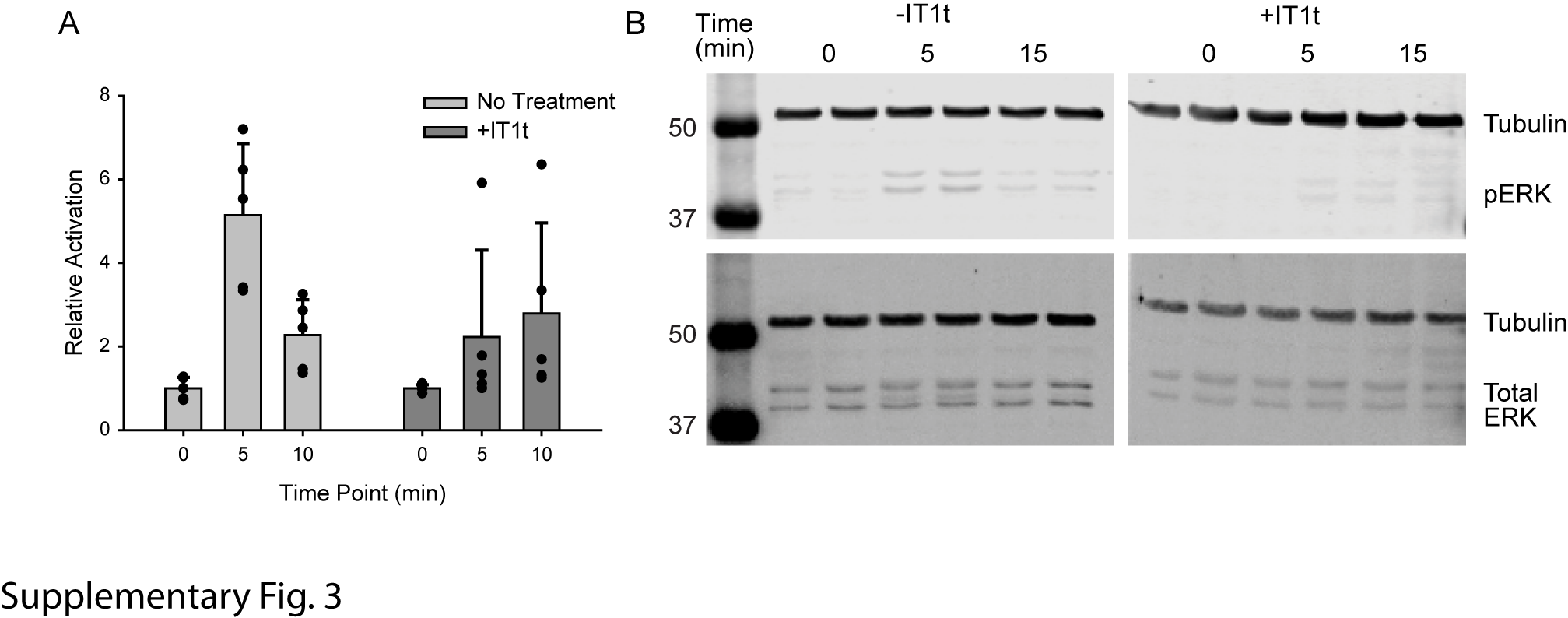

Supplementary Figure 3: CXCL12-mediated ERK phosphorylation is due to CXCR4 activation in our HEK293A cells and not ACKR3 activation. A) Relative activation of phospho-ERK reported as a fraction of total ERK and normalized to the ratio at t=0 in the absence or presence of 100 µM IT1t. Experiments were conducted in quadruplicate and error bars reflect standard deviations. B) Western blots reporting phospho-ERK (top) and total ERK (bottom) used to determine the ratios presented in panel A.
